# The evolution of sexual dichromatism in a large radiation of landfowl: Re-examining female-biased selection in Wallace’s model

**DOI:** 10.1101/2022.01.02.474725

**Authors:** Zheng Li, De Chen, Lu Dong

## Abstract

Sexual dichromatism is generally thought to arise from sexual selection favouring elaborately coloured males, as proposed in Darwin’s model. Wallace offers an expanded perspective, emphasising that the evolution of cryptic and dim female colouration to evade nest predation significantly contributed to the origin of sexual dichromatism. However, previous studies on the evolutionary forces of sexual dichromatism examining Darwin’s and Wallace’s models have produced mixed results. Here, we re-examined Wallace’s model of female-biased selection using the largest and most ecologically diverse family of landfowl (Phasianidae), known for its wide-ranging distribution and striking plumage patterns. Our results demonstrate that the level of sexual dichromatism is negatively correlated with colour contrasts in females but not males, and the evolutionary rates of sexual dichromatism are positively correlated with the evolutionary rates of colour contrasts in both sexes. Furthermore, we show that species with lower nest safety have significantly reduced female colour contrasts and sexual dichromatism, supporting Wallace’s model by highlighting the role of nest predation in driving evolutionary changes in sexual dichromatism within the Phasianidae family.

## Introduction

Sexual dichromatism, a complex characteristic that expresses the colour differences between male and female, has long enchanted evolutionary biologists, especially in the study of birds (Darwin, 1871; Cronin, 1993; Andersson, 1994). Sexual dichromatism is generally thought to be a result of increased ornamentation in males, as such bright and conspicuous colours are widely assumed to represent genuine sexual signals (Amundsen, 2000; Badyaev & Hill, 2003). However, dichromatism arose when the two sexes were under different selection, with increased or decreased ornamentation in either or both males and females (Kimball & Ligon, 1999; Amundsen, 2000; Badyaev & Hill, 2003). A growing number of studies have found that changes in female plumage colouration could drive the evolution of sexual dichromatism in different groups (Irwin, 1994; Price & Whalen, 2009; Johnson *et al*., 2013; Price & Eaton, 2014; Dale *et al*., 2015; Maia *et al*., 2016), but to date, the results explaining the contributions of females or males to sexual dichromatism have been mixed (Shultz & Burns, 2017; Cooney *et al*., 2019).

The debate over evolutionary causes of sexual dichromatism has long centred on two hypotheses. Darwin’s classic model holds that increased ornamentation of elaborate and conspicuous plumage in males, driven by sexual selection, is the sole explanation of the evolution of sexual dichromatism (Darwin, 1871). However, it fails to explain why females in sexually monomorphic species, such as bee-eaters, parroquets, and francolins, are as bright and colourful as males. Wallace resolved this contradiction by suggesting that selection due to nest predation drive the evolution of dichromatism, as females with drab and inconspicuous colouration are more likely to survive in open nests, while cavity or domed nesting females can exhibit more vibrant plumage under relaxed selection pressures (Wallace, 1868; Soler & Moreno, 2012). Although research on certain groups, including tanagers (Shultz & Burns, 2017) and suboscine passerines (Cooney *et al*., 2019), indicates that sexual dichromatism correlates with the evolution of male plumage colouration due to sexual selection, a growing number of studies suggests that dichromatism evolves through a mosaic of sexual and natural selection in both sexes (Shultz & Burns, 2017). Recently, multiple studies have increasingly demonstrated that the evolutionary gains of dichromatism are prominently due to losses in female elaboration, supported by empirical studies in passerines (e.g., in new world orioles (Friedman *et al*., 2009), new world blackbirds (Irwin, 1994), Australian fairy-wrens (Johnson *et al*., 2013), grackles-and-allies (Price & Eaton, 2014), starlings (Maia *et al*., 2016) and all passerines (Dale *et al*., 2015). However, previous studies have primarily focused on select avian groups with limited colour diversity, especially in passerine birds (Irwin, 1994; Burns, 1998; Johnson *et al*., 2013; Price & Eaton, 2014; Dale *et al*., 2015; Cooney *et al*., 2019), and have not thoroughly examined the link between selective forces and the evolution of colouration in both sexes.

Conspicuousness in birds can be expressed through various colour patterns with each tailored to meet specific ecological and social needs. Plumage brightness has often been used as an indicator of signal strength, with bright colours interpreted as an honest sexual signal (McGraw, 2006a, 2006b; Dale *et al*., 2015) and duller colours seen as a result of natural selection pressures (Wallace, 1889). However, Previous studies that used brightness to investigate the evolution of sexual dichromatism have not distinguished between similar brightness levels that differ significantly in colour composition (Dale *et al*., 2015). Additionally, conspicuousness can also result either from the contrast between an individual and its background or from contrasts among various plumage patches within the individual (Endler, 1990). By incorporating variations in hue and saturation, colour contrast (complexity) captures a broader range of visual signals than brightness alone, which is particularly relevant in species where males and females display distinct colours (Shultz & Burns, 2017). Research on manakins has shown that within-plumage colour contrasts is correlated with the individual colour contrast against background (Doucet *et al*., 2007). Therefore, measuring colour contrasts among patches within an individual can be a more effective way to study colour evolution in terms of conspicuousness, given the challenges of assessing background contrasts in diverse habitats.

The Phasianidae family (i.e., pheasants, quails, grouse and allies) provides an ideal system for this investigation due to its extreme variation in dichromatism levels and complex life histories potentially causing distinct selective forces. Phasianidae is a species-rich family with worldwide distribution and occupancy in nearly all terrestrial habitats from alpine tundra to tropical rainforest (Johnsgards, 1999). The various species within Phasianidae exhibit a wide diversity of plumage patterns ranging from highly contrasting colour patches to more cryptic, brown-toned plumages that blend into their surroundings. Many Phasianidae species, such as the red jungle fowl (*Gallus gallus*) and Indian peafowl (*Pavo cristatus*), are well-known for their sexual dichromatism and are thought to be under strong sexual selection pressures (Winkler *et al*., 2015), as proposed in Darwin’s model. However, in most Phasianidae species, females are solely responsible for incubation and other parental care duties, which exposes them to intense natural selection pressures from predation (Collias & Collias, 2014). This aligns with the expectations of Wallace’s model, where females may evolve cryptic colouration as an adaptation to minimize predation risk.

In this study, we conduct a large-scale comparative analysis of the evolution of sexual dichromatism, examining the relationship between dichromatism and the evolutionary trajectories of colour changes in both sexes in terms of direction and rates. By incorporating predictors that capture different selective pressures, we aim to explain the mechanisms driving these changes, offering a deeper understanding of the evolutionary dynamics influencing sex-biased colouration differences. We tested two predictions from Wallace’s model: (1) We first examined whether increased dichromatism is negatively correlated with reduced colour contrasts in females by relating levels of dichromatism and colour contrast across both sexes. (2) Using ecological factors including mating system, nest safety (including nest structure and nest sites), and habitat density, we tested whether decreased colour contrast in females is associated with increased nest predation risk. Conversely, we examined whether increased colour contrast in females correlates with decreased predation risk, represented by closed nests and lower nest sites.

## 2. Materials and methods

### (a) Colour quantification

We obtained images of Phasianidae species from the plates in the *Handbook of the Birds of the World* (HBW) (Del Hoyo *et al*., 1992) at https://birdsoftheworld.org. These images were read into the R statistical programming environment (R Core Team, 2020) for analysis. We measured the RGB (red, green, blue) values for eight patches (nape, crown, forehead, throat, upper breast, lower breast, face, and lore) on each sex of each species with the R package “colorZapper” (Valcu & Dale, 2014). We chose these regions because (1) they are very important signalling regions in Phasianidae showing a variety of colouration (Price-Waldman *et al*., 2020; Beltrán *et al*., 2021), and (2) they are illustrated clearly in the plates such that each patch of all species can be accurately measured. For each patch except the lore, a polygon was subjectively selected that contained the representative colouration in that area of the bird’s plumage (Figure 1a). We extracted RGB values using colorZapper for 400 optionally chosen pixels within the selected polygon. For lore, five points around the eye were randomly selected, representing the bare skin around the eye (Figure 1a), of which the RGB values were exported by colorZapper. We calculated the mean values for R, G, and B from the pixels or points for each patch, and then we were able to obtain a set of RGB values for each patch, separately for each sex within each species (Table S1). The deficiency of UV vision in Phasianidae species makes the manual image-based RGB measurement reasonable for colour analysis (Hart *et al*., 1999; Hart, 2002; Lind & Kelber, 2009).

**Figure 1.**
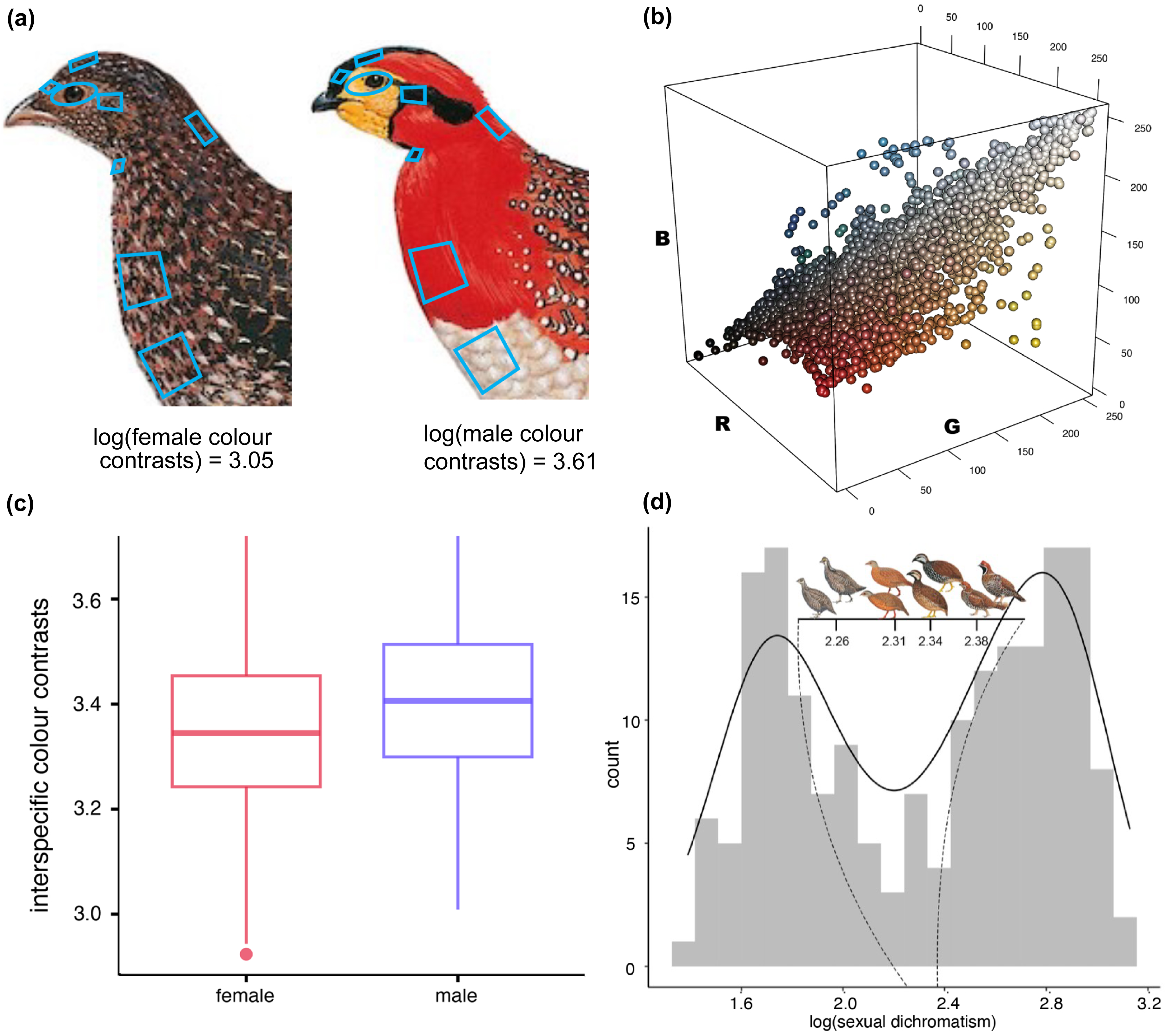
(a) Example of eight patches chosen for measuring RGB values in gray-bellied tragopan (*Tragopan blythii*): polygons selected on seven regions (nape, crown, forehead, throat, upper breast, lower breast, face), and five points selected around eyes (lore). (b) The colour of each patch of Phasianidae is shown in RGB space. For each patch, colour values for both sexes in all species were pooled (N = 1,464 points, where the point colour is determined from actual RGB values). (c) Boxplot of interspecific colour contrasts within all eight regions measured for all species for males and females. (d) Bimodal distribution of sexual dichromatism levels in Phasianidae. The dashed line interval and plates show the transition from sexual monochromatic species to sexual dichromatic species.

For those species with posted male and female images in HBW (N = 113), we used different images of both sexes to extract RGB values. However, as some monochromatic species (N = 70) have only one image showed in the HBW, we therefore sampled the same image twice independently, once for each sex. This processing was performed to maintain a similar measuring error between monochromatic species and dichromatic species.

We visualised RGB values in a three-dimensional colour space (Figure 1b). The scattered points from black (0, 0, 0) to white (255, 255, 255) form a diagonal line obliquely across the RGB colour space, and the rest of the scattered colour points spread around the diagonal line and form a curved surface (Figure S1).

### (b) Evaluation of colour accuracy

To evaluate whether the drawings from the HBW yield a valuable representation of the true colour of the Phasianidae species in nature, we utilised a digital photographic method to quantify the colour of museum specimens and compared these results to the colour of the drawings from the HBW.

We measured 30 specimens from 18 Phasianidae species at the Natural History Museum of Beijing Normal University, representing good coverage across both phylogeny and different levels of dichromatism (Table S2). Photos were taken with a Canon EOS 80D with a Canon EF 70-300 mm f/4-5.6L telephoto zoom lens. The specimen was illuminated with two 400 W photoflood lamps using a softbox to soften the light falling on the specimen. All photos were taken at an ISO of 100, with aperture of f/8.0 and 1/125 sec shutter speed, with the shooting distance being slightly adjusted so that the selected colour area could be fully displayed in each photo. Datacolor’s SpyderCHECKR 24 colour card was used to adjust for slight variation in separate light environments and standardise the colourations between photos in Adobe Lightroom. For these photos of specimens, we also quantified patch colour using the same procedure as for the drawings (Table S2). The R, G, and B values were found to correlate well between the drawings and specimen photos, with Pearson correlation coefficients (r) of 0.83, 0.80, and 0.79, respectively (*P* < 2.2×10^-6^, Figure S2). Although the R, G, and B values of the drawings were larger than the values of the photos (regression slopes were 0.68, 0.63, and 0.54), the absolute deviation does not affect our analyses since these differences were consistent with the phylogeny.

### (c) Phylogeny and divergence times

To obtain the timetree of all 183 species of Phasianidae, we used recently published genus-level topology for Galliformes from ultraconserved elements (UCEs) (Chen *et al*., 2021) as a backbone tree to obtain phylogenetic relationships for 159 Phasianidae species that have nucleotide sequences in GenBank (Table S3). We obtained CYTB, ND2, CLTC, CLTCL1, EEF2, FGB, RHO, SERPINB14, and OVOG from GenBank, as these genes are widely sequenced in Galliformes (Crowe *et al*., 2006; Wang *et al*., 2013). We added 12 representative species from the other four families in Galliformes, as these species have higher sequence coverage for the selected genes. Three outgroups, Common ostrich (*Struthio camelus*) and two ducks (*Anas platyrhynchos*, *Oxyura jamaicensis*), were used to provide more distant outgroups to reduce the stochastic error in time estimation (Wang *et al*., 2017). We used RAxML v8.2.12 (Stamatakis, 2014) under a best tree plus 100 rapid bootstrap replicates (‘-f a’ option) using the GTR + G model to reconstruct the maximum-likelihood tree.

Fixing the tree topology, we estimated the divergence times in MCMCTREE (PAML 4.9j) (Yang, 2007) with approximate likelihood calculation. Five carefully chosen galliform fossils that were validated (Hosner *et al*., 2016, 2017; Wang *et al*., 2017) were used as lower minimal bounds for node ages, and a maximum age constraint of 99.6 million years ago (Ma) for the tree root was set. The effective sample sizes of all parameters were checked in Tracer 1.7.1 (Rambaut *et al*., 2018) to ensure that they were above 200.

After obtaining the divergence times from the 159 Phasianidae species, we manually added the missing 24 Phasianidae species using TreeGraph 2 (Stöver & Müller, 2010). The node for each missing species was arbitrarily placed at the middle of the branch of its closest relative based on other phylogenetic studies in Phasianidae when possible (e.g., (Chen *et al*., 2020; Kimball *et al*., 2021)), or according to its morphological relatedness (data from HBW). This method has been widely used to integrate missing species into phylogeny and showed little effect in large-scale analyses (Algar *et al*., 2009; Cai *et al*., 2018).

### (d) Ancestral state reconstruction of sexual dichromatism and colour contrasts

We calculated the Euclidean distance of each patch using RGB values between the two sexes within species, the sum of which was taken as the level of sexual dichromatism of species (Table S4). The Colour contrasts from males and females were calculated by summing the Euclidian distances among all patches of each species (Table S4). To compress the scale of sexual dichromatism, we used log-transformed values in the statistical analysis. To infer the evolutionary history of the sexual dichromatism and colour contrasts along the phylogeny, we performed maximum likelihood ancestral state reconstructions, implemented with the R package “phytools” (Revell, 2012), which estimates the levels of sexual dichromatism and colour contrasts at ancestral nodes on the phylogeny of Phasianidae. The phylogenetic signal of dichromatism was tested using Pagel’s lambda (λ) statistics with the R package “phytools” (Revell, 2012).

### (e) Estimation of colour evolutionary rates

To calculate rates of phenotypic evolution, we applied a recently developed ridge regression method, implemented with the functions from R package “RRphylo” (Castiglione *et al*., 2018). RRphylo is designed to quantify phenotypic evolutionary rates and their variation across phylogenetic trees and locate rate shifts relating to entire clades, as well as unrelated tree tips. Compared to classic, model-based phylogenetic comparative methods, RRphylo has no priori assumptions concerning the tempo and mode of phenotypic evolution. However, rates are presumed to change faster among clades than within clades, and the rate of change is calculated as proportional to time (branch lengths). The sign and level of the regression slope at each branch estimated with RRphylo refer to the direction and magnitude of the phenotypic evolution, respectively.

The evolutionary rates of sexual dichromatism and colour contrast were both estimated by RRphylo. Here, we used the log-transformed magnitude representing rates for statistical analysis because the statistical analysis of the direction of phenotypic evolution was performed independently.

### (f) Comparative analyses

To explore the relationship between male and female colour contrasts and sexual dichromatism, we first examined whether higher within-body contrasts in female or male colourations associated with greater levels of sexual dichromatism. Subsequently, we assessed whether lineages with higher evolutionary rates of sexual dichromatism were related to increased evolutionary rates of male or female colour contrasts, using rates estimated with RRphylo. We fit phylogenetic generalised least squares (PGLS; (Grafen, 1989; Martins & Hansen, 1997)) models to analyze the relationships between: (1) male colour contrasts and sexual dichromatism, (2) female colour contrasts and sexual dichromatism, (3) evolutionary rates of male colour contrasts (RMC) and evolutionary rates of sexual dichromatism (RSD), and (4) evolutionary rates of female colour contrasts (RFC) and RSD. These analyses were conducted using functions from the R packages “ape” (Paradis *et al*., 2004) and “nlme” (Pinheiro *et al*., 2019), implementing Pagel’s model of continuous character evolution (Pagel, 1999).

To further test the relative contribution of male and female colour contrasts to the gains and losses of sexual dichromatism, we categorised the RSD as increased (positive rates) or decreased (negative rates). We then classified species into groups exhibiting evolutionary increased sexual dichromatism (iSD) or decreased sexual dichromatism (dSD) according to these categories.

To investigate the evolutionary forces influencing colour contrasts in both sexes and their role in the evolution of sexual dichromatism, PGLS (Grafen, 1989; Martins & Hansen, 1997) was used to assess the effects of ecological and life history traits on sexual dichromatism and colour contrasts. Specifically, we included mating systems (scored as monogamous [0] or polygamous [1]), nest safety (from 1 to 4 following Delhey et al., 2023), and habitat density (scored as dense [1], semi-open [2] or open [3]). Nest safety scores were derived by combining nest site scores (1 for ground nests, 2 for nests in vegetation, 3 for nests on cliffs or banks) and nest structure scores (0 for closed nests, 1 for open nests), with intermediate scores assigned to species using multiple nest types. Data were collected from published sources, with detailed data references provided in Table S5. The dataset for these three predictors included 173 species, representing 95% of all analysed species (Table S4). Model performance was assessed using Akaike Information Criterion (AIC), and effect sizes (β) were interpreted to infer the relative importance of predictors.

## 3. Results

The distribution of sexual dichromatism across Phasianidae species showed a distinct bimodal pattern (Figure 1d), with strong phylogenetic signals (Pagel’s λ = 0.91, *P* = 1.1×10^-28^) and considerable variation among clades. The interspecific colour contrasts were significantly higher in males than females across all species pairs (paired t-test: t = 4.14, df = 182, *P* < 0.01; Figure 1c), indicating greater colour variety in males.

Ancestral state reconstruction reveals that the Phasianidae ancestor likely displayed moderate sexual dichromatism (Figure 2c). The reconstructed ancestral states of colour contrasts demonstrated a complex evolutionary pattern where diverse evolutionary processes between males and females interact, with multiple independent gains and losses of sexual dichromatism across the phylogeny (Figure 2a, b).

**Figure 2.**
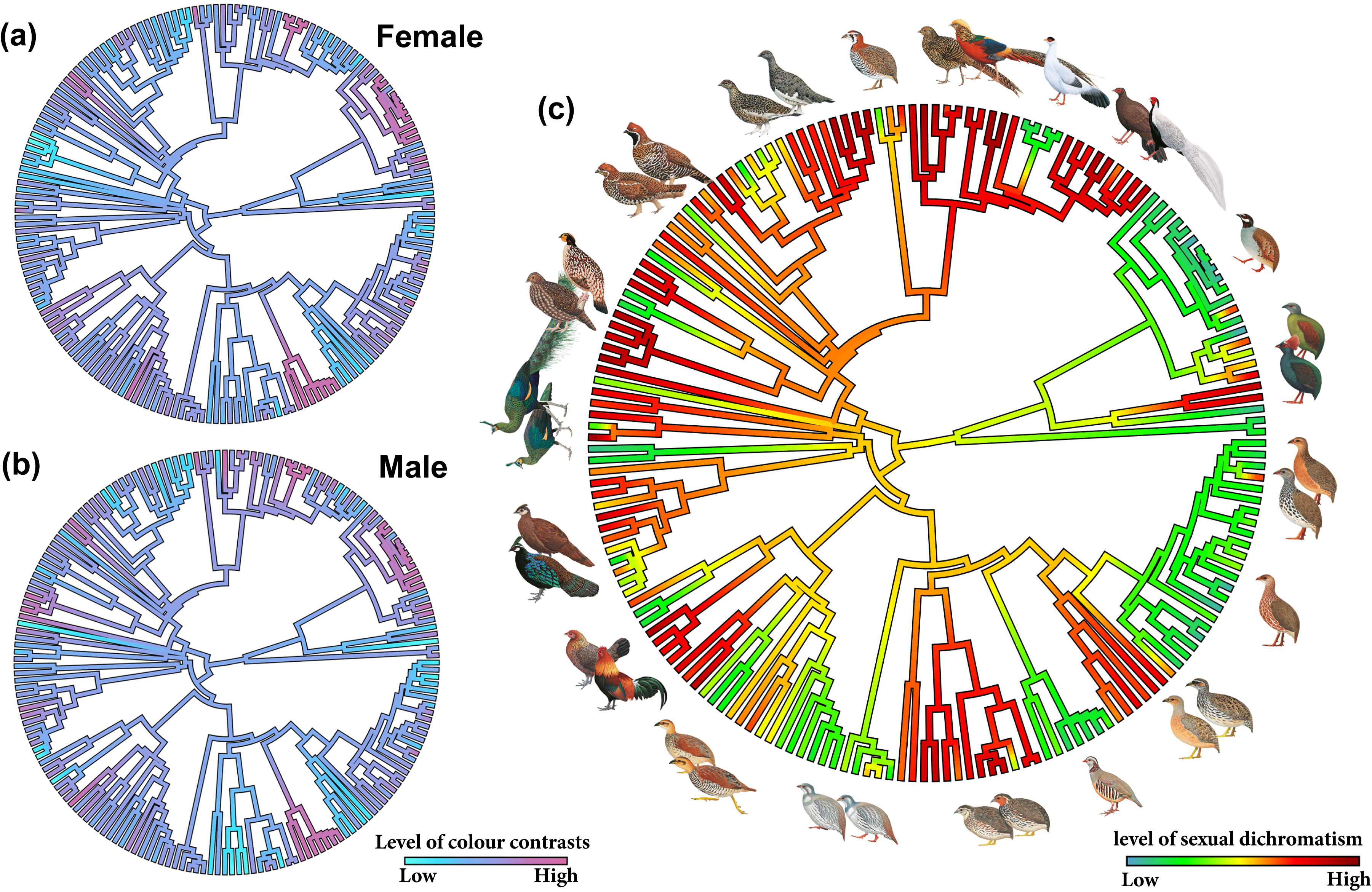
Evolutionary reconstructions of (a) colour contrasts in males, (b) females and (c) sexual dichromatism in Phasianidae in the molecular phylogeny. Colours mapped on the branches represent the level of colour contrasts (blue: low; pink: high) in (a, b) and sexual dichromatism (green: low; red: high) in (c).

Analysis of evolutionary rates showed no significant trend in sexual dichromatism (t = −0.724, df = 304.71, *P* = 0.47, Figure S3c) or male colour contrasts (t = 1.527, df = 333.51, *P* = 0.13, Figure S3a) across the clade. However, female showed a significant accelerating trend (t = 3.684, df = 218.07, *P* < 0.01) as colour contrasts increase, suggesting a unique rapid shift in female colour evolution (Figure S3b).

Sexual dichromatism was negatively correlated with female colour contrast (β = −0.85, *P* < 0.01, Figure 3b) but not with male colour contrast (Figure 3a). The analyses of sexual dichromatism, categorised into increased and decreased stages, revealed negative correlations with female colour contrasts in both categories (iSD: R = −0.36, *P* < 0.01, dSD: R = −0.47, *P* < 0.01; Figure 4a,b) whereas no such correlations were observed in males (Figure 4a,b).

**Figure 3.**
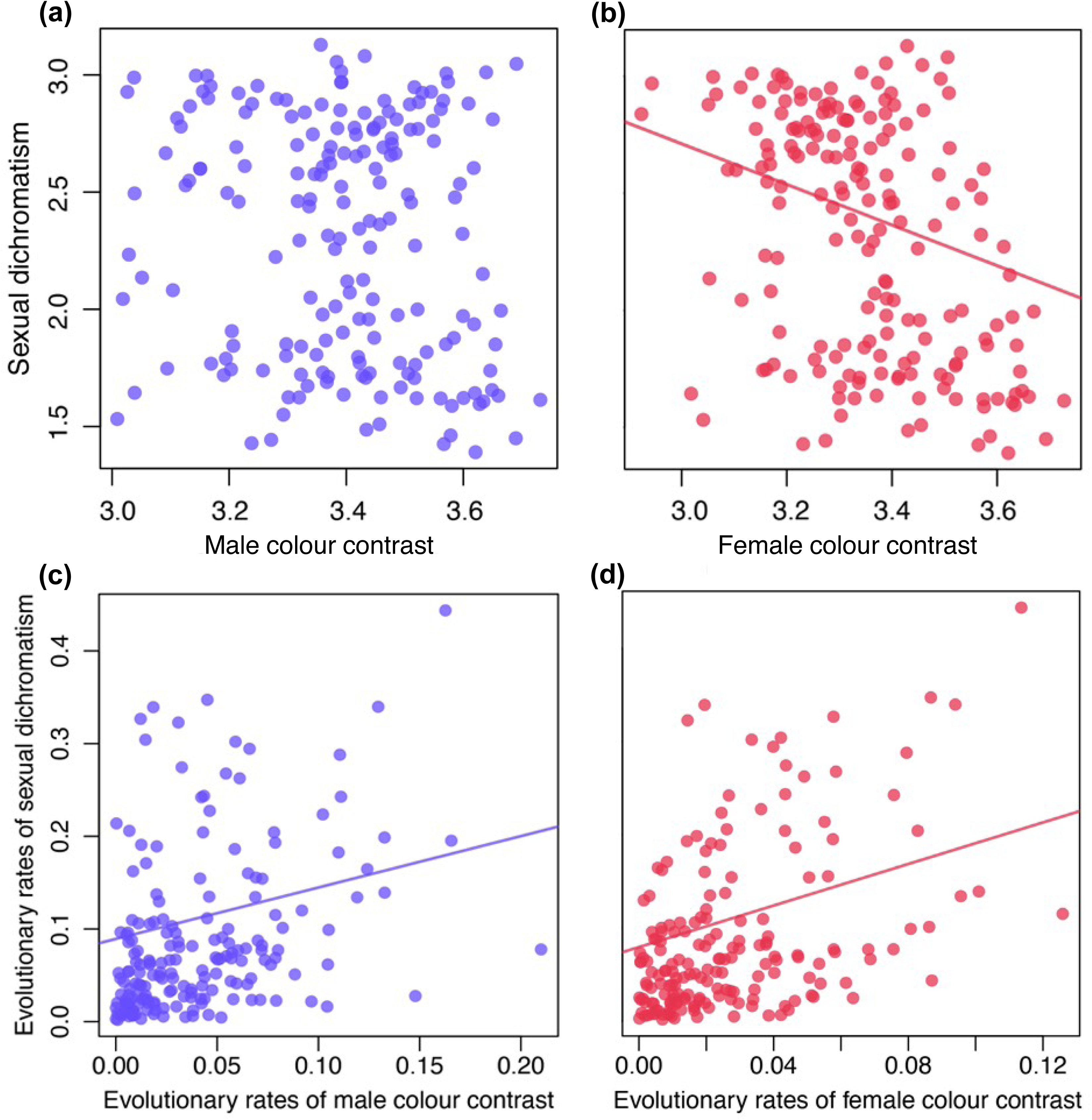
Relationships between colour contrasts and dichromatism estimated by PGLS regression (red: female; blue: male). (a) Male colour contrasts and sexual dichromatism, (b) female colour contrasts and sexual dichromatism (β = −0.85, P < 0.0001), (c) evolutionary rates of male colour contrasts and evolutionary rates of sexual dichromatism (β = 0.55, P < 0.001), (d) evolutionary rates of female colour contrasts and evolutionary rates of sexual dichromatism (β = 1.11, P < 0.0001).

**Figure 4.**
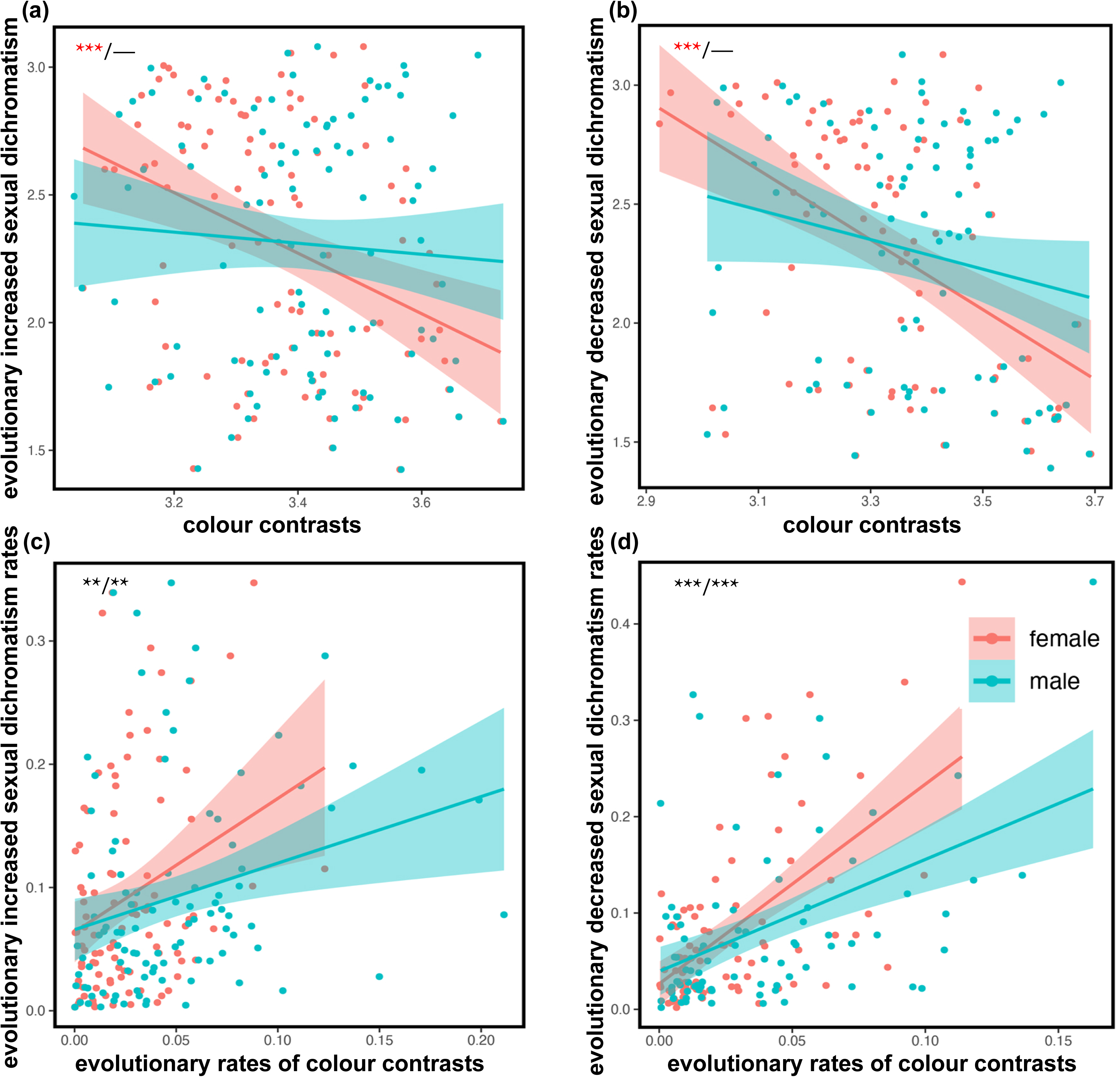
Correlations between colour contrasts and the processes of evolutionary increased and decreased sexual dichromatism. (a) Evolutionary increased sexual dichromatism and colour contrasts, (b) evolutionary decreased sexual dichromatism and colour contrasts, (c) evolutionary increased rates of sexual dichromatism and rates of colour contrasts, (d) evolutionary increased rates of sexual dichromatism and rates of colour contrasts. Regression lines are fitted in a Pearson linear model. Asterisks indicate significance, where ** represents *P* < 0.01 and *** represents *P* < 0.001 (red: negative correlation, black: positive correlation).

The evolutionary rates of sexual dichromatism showed positive correlation with evolutionary rates of colour contrasts in both females (β = 1.11, *P* < 0.0001, Figure 3d) and males (β = 0.55, *P* < 0.001, Figure 3c). Notably, females exhibited faster evolution than males along branches with marked changes in sexual dichromatism, indicating that the dichromatism evolves align with colour contrast changes across both sexes, with females showing more directional evolution. The evolutionary increased and decreased sexual dichromatism rates were both positively correlated with RFC (iRSD: R = 0.29, *P* < 0.01, dRSD: R = 0.6, *P* < 0.001; Figure 4c,d) and RMC (iRSD: R = 0.27, *P* < 0.01, dRSD: R = 0.47, *P* < 0.001; Figure 4c,d), further supporting the influence of both male and female colour evolution on dichromatism shifts.

Model selection analyses revealed that nest safety scores were the most significant predictor of variation in sexual dichromatism, with more exposed nests positively correlated with higher sexual dichromatism (β = 0.10, *P* = 0.0132, Table S6). Models fitting female colour contrasts, based on mating system alone (AIC = −49.35342, Loglik = 27.67671) or in combination with nest safety scores (AIC = −46.39646, Loglik = 27.19823) exhibited similar performance. Both models demonstrated negative correlation between the polygamous mating system and female colour contrasts (mating system alone: β = −0.09, *P* < 0.05; mating system and nest safety scores: β = −0.10, *P* < 0.05). The combination model also revealed a negative correlation between exposed nests and female colour contrasts (β = −0.05, P < 0.05). We found that none of the examined predictors (including mating system, nest safety, or habitat density) showed significant relationship with male colour contrasts.

## 4. Discussion

Our large-scale comparative analysis of colour evolution in Phasianidae provides novel insights into the evolutionary dynamics of sexual dichromatism and sex-specific colour patterns. The results reveal a complex interplay of selective pressures shaping colour patterns of this diverse avian clade. Our results also showed that (i) sexual dichromatism is negatively correlated with colour contrasts in females but not in males. (ii) the evolutionary rates of dichromatism is positively correlated with the rates of colour contrasts in both sexes while females evolve faster where dichromatism is changing remarkably. (iii) dichromatism and less contrasting female colours are associated with exposed nests, while less contrasting female colours is also associated with polygamy. These findings collectively support the hypothesis that natural selection, particularly nest predation pressure, constrains female colouration, and reflect the interplay between natural and sexual selection, with rapid changes in female colouration driving shifts in overall dichromatism levels.

The observed bimodal distribution of dichromatism across Phasianidae species demonstrates the suitability of this clade as an ideal system to investigate the evolution of sexual dichromatism. Ancestral state reconstruction indicates that moderate sexual dichromatism was likely present in the common ancestor of Phasianidae, suggesting deep evolutionary origins. Furthermore, the recurrent gains and losses of dichromatism across the clade emphasize the importance of incorporating phylogenetic history when interpreting the present patterns of sexual dichromatism.

The negative correlation between dichromatism and female colour contrast (β = −0.85, *P* < 0.01, Figure 3b), observed consistently in both the increasing and decreasing stages of dichromatism (iSD: R = −0.36, *P* < 0.01, dSD: R = −0.47, *P* < 0.01; Figure 4a,b), suggests that evolutionary shifts in female colouration play a crucial role in both the gains and losses of dichromatism. As dichromatism increases, female contrasts tends to decreases, resulting in more cryptic and less conspicuous colour. This pattern reflects an adaptive strategy for reducing visibility to avoid predation (Irwin, 1994; Gomez & Théry, 2007), particularly in species with open nesting behaviors, as initially proposed by Wallace (Wallace, 1868). Comparative phylogenetic analysis further supports this hypothesis, showing a correlation between increased dichromatism, exposed nesting, and less contrasting female colours (Table S6). Furthermore, our observation that polygamous females tend to have lower colour contrasts supports the hypothesis that females often face higher predation pressures than males when they are responsible for incubation and parental care alone (Martin & Badyaev, 1996; Scarnecchia & Johnsgard, 2000; Palmer & Carroll, 2003).

Interestingly, we found that females exhibit an acceleration in colour contrasts as they increase (t = 3.684, df = 218.07, *P* < 0.01), suggesting that under intense natural selection, female colouration may be suppressed. However, in the absence of such pressures, relaxed natural selection may allow for less constraint on the evolution of female colouration, enabling rapid and unpredictable changes, which could potentially lead to a loss of sexual dichromatism. The observed acceleration in female color contrasts aligns is consistent with Price et al. (2024), which indicated that decreases in sexual dichromatism often involve rapid changes in female plumage colouration.

Conversely, male colours exhibit greater diversity and variation in interspecific contrasts than female colours (paired t-test: t = 4.14, df = 182, *P* < 0.001; Figure 1c). The absence of a correlation between male colour contrasts and sexual dichromatism suggests that female colour evolution follows a more defined direction and trend, while male colouration evolves along a more variable and stochastic path (Johnson *et al*., 2013; Price & Eaton, 2014). This divergence is likely driven by the differential interplay of sexual and natural selection on male and female colours. Despite these differing selective pressures, the positive correlation between the rates of colour contrast and dichromatism evolution in both sexes indicates that the broader evolutionary dynamics driving dichromatism influence colour evolution in parallel (Figure 3c,d). This suggests that while males and females evolve different colour traits due to sex-specific selective pressures, the rates of change in colour and dichromatism are linked through shared ecological and evolutionary contexts. This suggests that both sexual and natural selection play important roles in shaping sexual dichromatism, with distinct mechanism of colour evolution between the sexes (Dunn *et al*., 2015; Delhey & Peters, 2017; Shultz & Burns, 2017).

The contrasting evolutionary trajectories of male and female colouration challenge the traditional perspectives on the role of sexual dichromatism in signaling and mate choice. Although sexual dichromatism has long been employed as a proxy for sexual selection (Darwin, 1871; Cooney *et al*., 2019), recent studies across different animal groups have suggested that the diversity of sexual dichromatism is shaped by a complex interplay of both natural and sexual selection (e.g. in frogs (Bell & Zamudio, 2012) and hummingbirds (Beltrán *et al*., 2021)). Our findings in the large radiation of landfowl provide further evidence of this mosaic effect, indicating that sexual dichromatism in this group cannot be fully explained by sexual selection but is also influenced by ecological and survival pressures.

Additionally, research in birds has proposed a trade-off between the elaboration of visual ornaments, such as plumage dichromatism, and the diversification of acoustic mating signals, underscoring the importance of considering multiple signaling modalities when evaluating the role of sexual selection in phenotypic diversification (Cooney, 2018). These insights highlight the need for a more integrative approach to understanding sexual dichromatism, one that considers both the ecological context and the evolutionary processes underlying colour evolution in both sexes.

## Conclusion

In conclusion, our study of the large species-rich family of landfowl provides valuable insights into Wallace’s model for the evolution of sexual dichromatism. We demonstrate that the directional evolution of female colouration is crucial in the evolution of sexual dichromatism, with rapid increases in female colour contrast contributing to the loss of dichromatism. The correlations between dichromatism, female colour contrasts, and nest safety scores provide further support for Wallace’s model, demonstrating how nest predation influenced changes in female colouration and, consequently, dichromatism. Additionally, we show that the evolutionary rate of sexual dichromatism is associated with the rate of colour contrast in both sexes, suggesting that dichromatism is shaped by the complex interplay of natural and sexual selection. Overall, the findings of this species-level phylogenetic comparative analysis of pheasants and allies emphasise the need for further investigation into other avian lineages to better understand how sexual dichromatism evolves in birds, considering the joint influence of natural and sexual selection.

## Data accessibility

Data will be provided in the electronic supplementary material after accepted.

## Authors’ contributions

L.D. and Z.L. conceived the study; Z.L. and D.C. performed the study and analysed data; Z.L., D.C. and L.D. drafted the manuscript. All authors approved the final version of the manuscript.

## Supporting information

Supplementary meterial

## Acknowledgements

We thank the Natural History Museum of Beijing Normal University for access to specimens and Ning Wang, Sen Dong, and Guocheng Yang for their help in specimen photography. We thank Zihan Hong for the help in ecological data collection. We also thank Nan Lyu and Zitan Song for comments on an earlier version of this manuscript.

## References

Algar, A.C., Kerr, J.T. & Currie, D.J. (2009) Evolutionary constraints on regional faunas: whom, but not how many. Ecology letters, 12, 57–65.

Amundsen, T. (2000) Why are female birds ornamented? Trends in Ecology and Evolution.

Andersson, M. (1994) Sexual selection. Princeton Univ. Press, Princeton.

Badyaev, A. V. & Hill, G.E. (2003) Avian Sexual Dichromatism in Relation to Phylogeny and Ecology. Annual Review of Ecology, Evolution, and Systematics, 34, 27–49.

Bell, R.C. & Zamudio, K.R. (2012) Sexual dichromatism in frogs: natural selection, sexual selection and unexpected diversity. Proceedings of the Royal Society B: Biological Sciences, 279, 4687–4693.

Beltrán, D.F., Shultz, A.J. & Parra, J.L. (2021) Speciation rates are positively correlated with the rate of plumage color evolution in hummingbirds. Evolution, 75, 1665–1680.

Burns, K.J. (1998) A phylogenetic perspective on the evolution of sexual dichromatism in tanagers (Thraupidae): The role of female versus male plumage. Evolution, 52, 1219– 1224.

Cai, T., Fjeldså, J., Wu, Y., Shao, S., Chen, Y., Quan, Q., et al. (2018) What makes the Sino-Himalayan mountains the major diversity hotspots for pheasants? Journal of Biogeography, 45, 640–651.

Castiglione, S., Tesone, G., Piccolo, M., Melchionna, M., Mondanaro, A., Serio, C., et al. (2018) A new method for testing evolutionary rate variation and shifts in phenotypic evolution. Methods in Ecology and Evolution, 9, 974–983.

Chen, D., Hosner, P.A., Dittmann, D.L., O’Neill, J.P., Birks, S.M., Braun, E.L., et al. (2021) Divergence time estimation of Galliformes based on the best gene shopping scheme of ultraconserved elements. BMC ecology and evolution, 21, 1–15.

Chen, D., Liu, Y., Davison, G., Yong, D.L., Gao, S., Hu, J., et al. (2020) Disentangling the evolutionary history and biogeography of hill partridges (Phasianidae, *Arborophila*) from low coverage shotgun sequences. Molecular phylogenetics and evolution, 151, 106895.

Collias, N.E. & Collias, E.C. (2014) Nest building and bird behavior. Princeton University Press.

Cooney, C.R., Varley, Z.K., Nouri, L.O., Moody, C.J.A., Jardine, M.D. & Thomas, G.H. (2019) Sexual selection predicts the rate and direction of colour divergence in a large avian radiation. Nature Communications, 10, 1–9.

Cronin, H. (1993) The ant and the peacock: Altruism and sexual selection from Darwin to today. Cambridge University Press.

Crowe, T.M., Bowie, R.C.K., Bloomer, P., Mandiwana, T.G., Hedderson, T.A.J., Randi, E., et al. (2006) Phylogenetics, biogeography and classification of, and character evolution in, gamebirds (Aves: Galliformes): effects of character exclusion, data partitioning and missing data. Cladistics, 22, 495–532.

Dale, J., Dey, C.J., Delhey, K., Kempenaers, B. & Valcu, M. (2015) The effects of life history and sexual selection on male and female plumage colouration. Nature, 527, 367–370.

Darwin. (1871) The Descent of Man, and Selection in relation to Sex. The Descent of Man, and Selection in relation to Sex.

Delhey, K. & Peters, A. (2017) The effect of colour-producing mechanisms on plumage sexual dichromatism in passerines and parrots. Functional Ecology, 31, 903–914.

Delhey, K., Valcu, M., Muck, C., Dale, J. & Kempenaers, B. (2023) Evolutionary predictors of the specific colors of birds. Proceedings of the National Academy of Sciences, 120, e2217692120.

Doucet, S.M., Mennill, D.J. & Hill, G.E. (2007) The evolution of signal design in manakin plumage ornaments. American Naturalist, 169, S62–S80.

Dunn, P.O., Armenta, J.K. & Whittingham, L.A. (2015) Natural and sexual selection act on different axes of variation in avian plumage color. Science advances, 1, e1400155.

Endler, J.A. (1990) On the measurement and classification of colour in studies of animal colour patterns. Biological Journal of the Linnean Society, 41, 315–352.

Friedman, N.R., Hofmann, C.M., Kondo, B. & Omland, K.E. (2009) Correlated evolution of migration and sexual dichromatism in the new world orioles (ICTERUS). Evolution, 63, 3269–3274.

Gomez, D. & Théry, M. (2007) Simultaneous crypsis and conspicuousness in color patterns: Comparative analysis of a neotropical rainforest bird community. American Naturalist, 169, S42–S61.

Grafen, A. (1989) The phylogenetic regression. Philosophical transactions of the Royal Society of London. Series B, Biological sciences, 326, 119–157.

Hart, N.S. (2002) Vision in the peafowl (Aves: P*avo cristatus*). Journal of Experimental Biology, 205, 3925–3935.

Hart, N.S., Partridge, J.C. & Cuthill, I.C. (1999) Visual pigments, cone oil droplets, ocular media and predicted spectral sensitivity in the domestic turkey (*Meleagris gallopavo*). Vision research, 39, 3321–3328.

Hosner, P.A., Braun, E.L. & Kimball, R.T. (2016) Rapid and recent diversification of curassows, guans, and chachalacas (Galliformes: Cracidae) out of Mesoamerica: Phylogeny inferred from mitochondrial, intron, and ultraconserved element sequences. Molecular Phylogenetics and Evolution, 102, 320–330.

Hosner, P.A., Tobias, J.A., Braun, E.L. & Kimball, R.T. (2017) How do seemingly non-vagile clades accomplish trans-marine dispersal? Trait and dispersal evolution in the landfowl (Aves: Galliformes). Proceedings of the Royal Society B: Biological Sciences, 284, 20170210.

Hoyo, J. Del, Hoyo, J. Del, Elliott, A. & Sargatal, J. (1992) Handbook of the birds of the world. Lynx edicions Barcelona.

Irwin, R.E. (1994) The evolution of plumage dichromatism in the New World blackbirds: Social selection on female brightness? American Naturalist, 144, 890–907.

Johnsgards, P.A. (1999) The pheasants of the world. Biology and Natural History.

Johnson, A.E., Jordan Price, J. & Pruett-Jones, S. (2013) Different modes of evolution in males and females generate dichromatism in fairy-wrens (Maluridae). Ecology and Evolution, 3, 3030–3046.

Kimball, R.T., Hosner, P.A. & Braun, E.L. (2021) A phylogenomic supermatrix of Galliformes (Landfowl) reveals biased branch lengths. Molecular Phylogenetics and Evolution, 158, 107091.

Kimball, R.T. & Ligon, J.D. (1999) Evolution of avian plumage dichromatism from a proximate perspective. American Naturalist, 154, 182–193.

Lind, O. & Kelber, A. (2009) Avian colour vision: effects of variation in receptor sensitivity and noise data on model predictions as compared to behavioural results. Vision research, 49, 1939–1947.

Maia, R., Rubenstein, D.R. & Shawkey, M.D. (2016) Selection, constraint, and the evolution of coloration in African starlings. Evolution, 70, 1064–1079.

Martin, T.E. & Badyaev, A. V. (1996) Sexual dichromatism in birds: Importance of nest predation and nest location for females versus males. Evolution, 50, 2454–2460.

Martins, E.P. & Hansen, T.F. (1997) Phylogenies and the comparative method: A general approach to incorporating phylogenetic information into the analysis of interspecific data. American Naturalist, 149, 646–667.

McGraw, K.J. (2006a) Bird Coloration: Mechanisms and Measurements. Mechanisms of melanin-based coloration.

McGraw, K.J. (2006b) Mechanics of melanin-based coloration. Bird coloration, 1, 243–294.

Pagel, M. (1999) Inferring the historical patterns of biological evolution. Nature.

Palmer, W.E. & Carroll, J.P. (2003) Pheasants, Partridges, and Grouse: A Guide to the Pheasants, Partridges, Quails, Grouse, Guineafowl, Buttonquails, and Sandgrouse of the World. The Auk, 120.

Paradis, E., Claude, J. & Strimmer, K. (2004) APE: Analyses of phylogenetics and evolution in R language. Bioinformatics, 20, 289–290.

Pinheiro, J., Bates, D., DebRoy, S., Sarkar, D. & R Core Team. (2019) nlme: Linear and nonlinear mixed effects models. https://cran.r-project.org/package=nlme.R-project.

Price, J.J. & Eaton, M.D. (2014) Reconstructing the evolution of sexual dichromatism: current color diversity does not reflect past rates of male and female change. Evolution, 68, 2026–2037.

Price, J.J., Garcia, K. & Eaton, M.D. (2024) Losses of sexual dichromatism involve rapid changes in female plumage colors to match males in New World blackbirds. Evolution, 78, 188–194.

Price, J.J. & Whalen, L.M. (2009) Plumage evolution in the oropendolas and caciques: Different divergence rates in polygynous and monogamous taxa. Evolution, 63, 2985– 2998.

Price-Waldman, R.M., Shultz, A.J. & Burns, K.J. (2020) Speciation rates are correlated with changes in plumage color complexity in the largest family of songbirds. Evolution, 74, 1155–1169.

R Core Team. (2020) R: a language and environment for statistical computing. See http://www.R-project.org/. Vienna, Austria: R Foundation for Statistical Computing.

Rambaut, A., Drummond, A.J., Xie, D., Baele, G. & Suchard, M.A. (2018) Posterior summarization in Bayesian phylogenetics using Tracer 1.7. Systematic biology, 67, 901.

Revell, L.J. (2012) phytools: An R package for phylogenetic comparative biology (and other things). Methods in Ecology and Evolution, 3, 217–223.

Scarnecchia, D.L. & Johnsgard, P.A. (2000) The Pheasants of the World. Biology and Natural History. Journal of Range Management, 53.

Shultz, A.J. & Burns, K.J. (2017) The role of sexual and natural selection in shaping patterns of sexual dichromatism in the largest family of songbirds (Aves: Thraupidae). Evolution, 71, 1061–1074.

Soler, J.J. & Moreno, J. (2012) Evolution of sexual dichromatism in relation to nesting habits in European passerines: A test of Wallace’s hypothesis. Journal of Evolutionary Biology, 25, 1614–1622.

Stamatakis, A. (2014) RAxML version 8: a tool for phylogenetic analysis and post-analysis of large phylogenies. Bioinformatics, 30, 1312–1313.

Stöver, B.C. & Müller, K.F. (2010) TreeGraph 2: combining and visualizing evidence from different phylogenetic analyses. BMC bioinformatics, 11, 1–9.

Valcu, M. & Dale, J. (2014) colorZapper: color extraction utilities. R package version 1.0.

Wallace, A.R. (1868) A theory of birds’ nests: showing the relation of certain sexual differences of colour in birds to their mode of nidification. J. Travel Nat. Hist, 1, 73–89.

Wallace, A.R. (1889) *Darwinism: an exposition of the theory of natural selection with some of its applications*. Macmillan, New York.

Wang, N., Kimball, R.T., Braun, E.L., Liang, B. & Zhang, Z. (2013) Assessing Phylogenetic Relationships among Galliformes: A Multigene Phylogeny with Expanded Taxon Sampling in Phasianidae. PLoS ONE, 8.

Wang, N., Kimball, R.T., Braun, E.L., Liang, B. & Zhang, Z. (2017) Ancestral range reconstruction of Galliformes: the effects of topology and taxon sampling. Journal of Biogeography, 44, 122–135.

Winkler, D.W., Billerman, S.M. & Lovette, I.J. (2015) Bird families of the world: an invitation to the spectacular diversity of birds. Lynx Edicions.

Yang, Z. (2007) PAML 4: phylogenetic analysis by maximum likelihood. Molecular biology and evolution, 24, 1586–1591.

