## Supplementary meterial for "The evolution of sexual dichromatism in a large radiation of landfowl: Re-examining female-biased selection in Wallace’s model"

Electronic Supplementary Material

Contents

Supplementary Figure

Gif plot of RGB values (Fig S1)

Correlations with RGB values in speciemens (Fig S2)

Evolutionary rates of sexual dichromatism and colour contrast (Fig S3)

External Database as an Excel file (Appendix S1)

**Supplementary** **Figure**

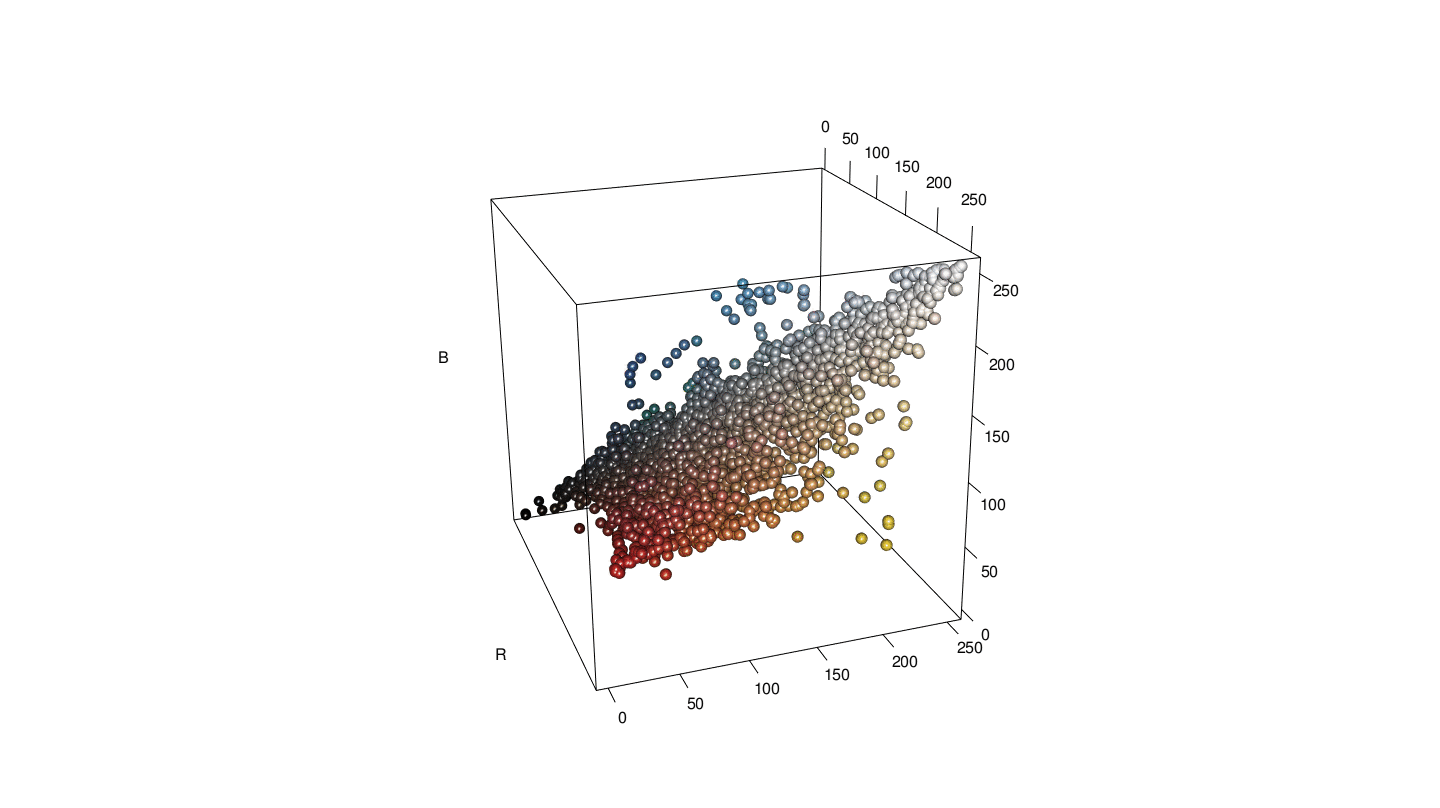

**Figure S1** The colour of each patch of Phasianidae shown in RGB space in gif version. For each patch, colour values for both sexes in all species were pooled (N = 1464 points, point colour is determined from actual RGB values). The X axis represents the values of R, the Y axis represents the values of G, the Z axis represents the values of B, and each point represents a RGB value of each patch from each sex of 183 species of Phasianidae.

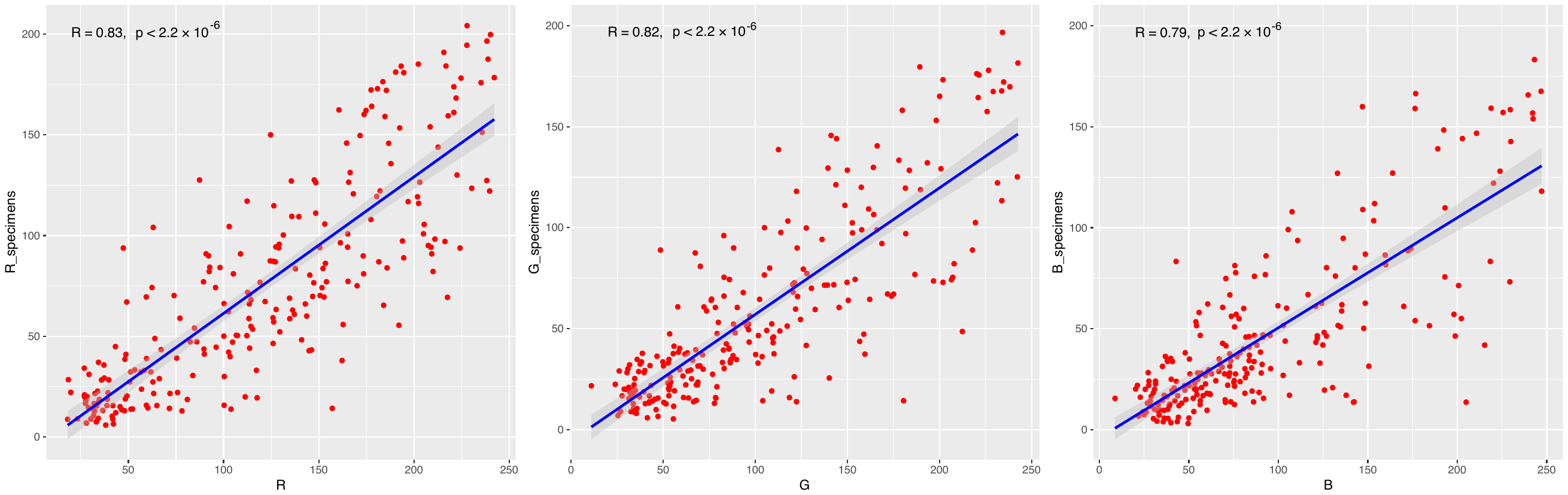

**Figure S2** The R, G, B correlation between drawings from the *Handbook of the Birds of the World* and 30 specimens collected from the Natural History Museum of Beijing Normal University.

**Figure S3** Evolutionary rates of (a) male colour contrasts, (b) female colour contrasts, (c) sexual dichromatism. Colours mapped on the branches represent absolute value of evolutionary rates calculated with RRphylo (cool colours = slow rates; warm colours = fast rates).
